# Intrinsic RNA hairpin-mediated transcription termination at high temperature in *Thermus aquaticus*

**DOI:** 10.64898/2025.12.25.696493

**Authors:** Nicholas P Bailey, Amber Riaz-Bradley, Yulia Yuzenkova

## Abstract

Transcription termination establishes gene boundaries and limits regulatory interference. In bacteria, intrinsic termination, mediated by an RNA hairpin followed by a polyuridine tract (polyU), is a principal mechanism. Because RNA secondary structures are destabilized at high temperature, whether intrinsic termination functions effectively in thermophiles has been uncertain. We mapped RNA 3′ termini genome-wide in the extreme thermophile *Thermus aquaticus* (*Taq*) grown at 70 °C using Term-Seq. Computational analysis identified hairpins upstream of most termini. These hairpins were substantially more thermally stable than those of the mesophile *Escherichia coli*. RNA 3′ ends were also detected within coding regions, consistent with transcriptional pausing or premature termination at spurious hairpins in GC-rich transcripts. *In vitro* transcription showed that native *Taq* terminators direct efficient intrinsic termination by *Taq* RNA polymerase at 70 °C and require both hairpin and polyU. Termination remained efficient up to 80 °C. At this temperature, the terminator hairpin RNA is unfolded in solution, indicating that RNA polymerase promotes hairpin formation during transcription. Thus, intrinsic termination operates efficiently in thermophiles. At lower temperatures, expression of the Rho termination factor increased, suggesting that factor-dependent termination may contribute more prominently under cooler growth conditions in *Thermus*.

**Significance Statement:** Intrinsic transcription termination requires formation of an RNA hairpin. Hairpin structures are expected to be unstable at the high temperatures inhabited by thermophilic bacteria. Whether and how hairpin-dependent termination operates under these conditions has remained unknown. Using genome-wide transcripts ends mapping and biochemical approaches, we show that the RNA polymerase of *Thermus aquaticus* promotes folding of the nascent RNA into a terminator hairpin. This enables efficient termination even at extreme temperatures. Raising temperature favours entropy-driven dissociation of the transcription complex. At lower temperatures, termination appears to increasingly rely on a factor-dependent pathway. Together, these results identify RNA polymerase as an active participant in hairpin-dependent termination and show how temperature may shift the balance between alternative termination mechanisms.

## INTRODUCTION

During transcription termination RNA polymerase (RNAP) stops and dissociates at defined sites, preventing readthrough into downstream loci and thereby preserving the independent regulation of adjacent genes. There are two mechanisms of transcription termination in bacteria, intrinsic termination (1–3), and via the termination factor Rho (4,5). Intrinsic terminators are defined nucleic acids sequence signals that cause RNAP to pause, destabilize the elongation complex, and release the nascent RNA without the involvement of dedicated termination factors. Elongation factors NusA and NusG influence termination efficiency for some bacterial RNAPs (6–8). Alternative mechanism of termination in bacteria requires an ATP dependent helicase Rho (9,10) which binds to specific rut sequences on the nascent RNA and translocates along the transcript until it disrupts elongation complex, leading to release of the RNA and termination of transcription. Rho may potentially in some cases assist intrinsic hairpin-dependent termination events (11).

A model intrinsic transcription terminator consists of a GC-rich palindromic sequence that forms an RNA hairpin followed immediately by a poly-uridine tract (polyU). The polyU promotes RNAP pausing providing time for RNA hairpin formation and contributes to destabilization of the elongation complex (2,12–15). A weak U–A base pairing reduces the stability of the DNA–RNA hybrid, facilitating collapse of the transcription bubble and release of the transcript once the hairpin has formed (16,17).

Three non-mutually exclusive models for elongation complex destabilisation during termination exist (18) - hypertranslocation (19) proposes that RNAP moves forward without adding nucleotides, shortening and destabilizing the RNA–DNA hybrid; strand slippage (20) suggests RNA hairpin mechanically disrupts the hybrid; and allosteric (21) where hairpin–RNAP interactions trigger conformational changes leading to RNA release.

Biochemical evidence supports contributions from all three mechanisms. Notably, hairpin formation plays a critical role in all models of transcription termination. Therefore, because intrinsic termination depends on formation of a weak RNA–DNA hybrid and a hairpin whose stability is inherently temperature sensitive, it is unclear how this mechanism operates in thermophilic bacteria. In hyperthermophilic archaea termination does not require RNA hairpin, and mostly depends on protein termination factors, yet can happen on a long homogeneous polyU sequence (22,23).

Indeed, RNA secondary structures necessary for RNA function at high temperatures require a high G+C content. A strong correlation has been observed between the G+C content of rRNA and tRNA - but not mRNA - and the optimal growth temperature across diverse prokaryotes, including *Thermus thermophilus* (*T. thermophilus*), suggesting that structural RNAs are thermally adapted (24). This raises the question - are intrinsic terminators hairpins also adapted for thermal stability?

*Taq* is a hyperthermophilic organism with a growth temperature range of 40-80 °C and optimum of 70 °C (25). To date, studies of intrinsic transcription termination in thermophilic eubacteria have been limited to moderate-temperature *in vitro* assays. In *T. thermophilus*, RNAP shows modest termination at 55°C under high NTP levels (26,27); higher termination at low NTPs (26,28). *T. thermophilus* NusA greatly enhanced cognate RNAP termination efficiency at 55°C (29), whereas NusG had a moderate positive effect on termination efficiency (27).

Intrinsic termination at physiological for hyperthermophiles temperatures (>60°C) has yet to be demonstrated *in vivo* and *in vitro*.

In this work, we asked whether elevated temperature constrains intrinsic transcription termination in *Thermus aquaticus* and whether and how native terminators are adapted to function at high temperature. Using Term-Seq specialised RNA-Seq technique which enriches transcript 3’ ends (30–32), we globally mapped transcript termini across the *T. aquaticus* genome and then characterized the determinants of termination efficiency *in vitro*. To assess the contribution of alternative termination mechanisms across the thermal range of *Thermus*, we further examined the temperature dependence of Rho factor expression.

## RESULTS

### 3’ RNA termini of *Taq* group into either sharp or “soft” peaks

Term-Seq analysis resulted in an average of 11.5 million sequence reads per sample across 3 biological replicates. Alignment to the reference genome predicted of 530 transcript 3’ termini which were concordant in all replicates. We defined 232 (44%) of these candidate terminators as “sharp” terminators, in which there was a large drop in Term-Seq read coverage over a 1 nt distance, indicating a defined 3’ terminus. 298 (56%) of the candidate terminators were defined as “soft” terminators, in which there was a more gradual decrease in read coverage, suggesting the presence of variable 3’ termini.

We re-analysed a published Term-Seq dataset from *E. coli* (32) with identical parameters, for use as a mesophilic control organism for comparison. We identified 595 transcript 3’ termini with concordant position across 3 experimental replicates. Of these, 571 (96%) were classified as “sharp” terminators.

### Features of hairpin and polyU elements of the *Taq* terminators

To assess potential intrinsic termination via the hairpin-polyU-dependent mechanism, we performed *in silico* RNA structure prediction on sequences surrounding the candidate terminators (50 bp upstream and 10 bp downstream of the predicted 3’ terminus), focusing on “sharp” terminators. Rescaling free energy from *in silico* RNA folding to a temperature of 70°C, 131 of the candidate 3’ termini (56% of the total “sharp” terminators) demonstrated predicted one or more hairpin with at least 4 paired bases and a predicted free energy of less than -10 kcal/mol. Two consecutive terminator hairpins were predicted for 35 of the terminators (15%). Seventy-eight (60% of termini with predicted hairpins) were located between 1-10 nt upstream of the predicted 3’ terminus (Figure 1A). The mean distance from the distal base of the hairpin at the termination site was 7 nt, consistent with Term-Seq defined termini from various bacterial species (33).

**Figure 1.**
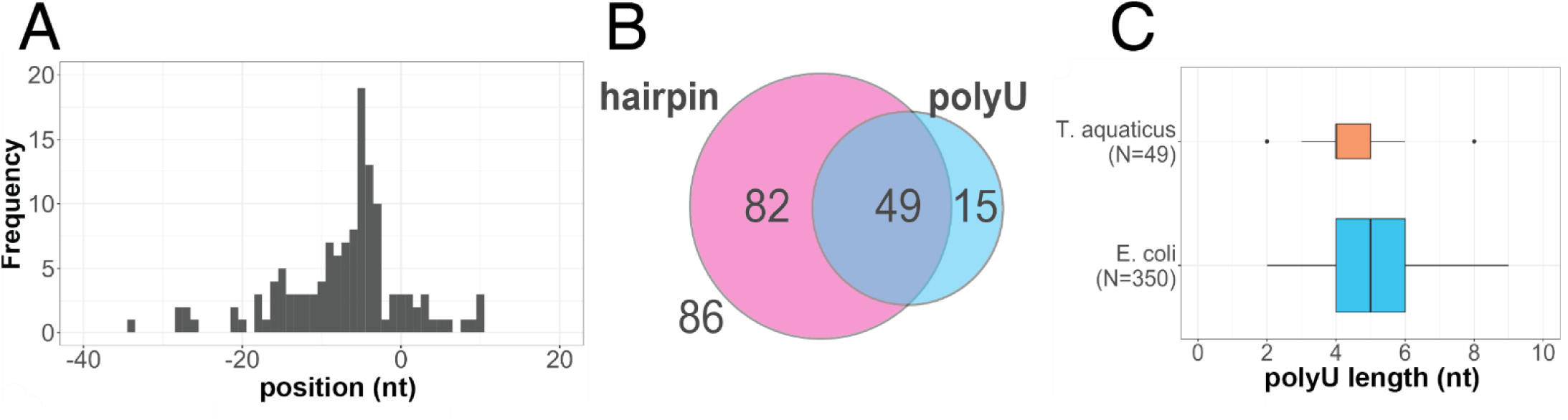
Intrinsic terminators in *Taq* predicted from Term-Seq data. **(A)** Frequency histogram showing the end position of *in silico* predicted hairpins relative to candidate “sharp” 3’ termini (at 0 nt position), including only termini proximal to a predicted hairpin. **(B)** Overlap of *in silico* predicted hairpin formation (rescaling free energy to 70°C) and polyU tract amongst “sharp” 3’ termini. **(C)** Comparison of polyU tract length proximal to the 3’ termini from *Taq* and *E. coli*, focusing only “sharp” termini for which both hairpins and polyU tracts were predicted according to the above criteria. Box height is scaled to square root of the sample size (which is indicated on the y-axis).

Amongst the termini with predicted hairpins, 49 included a polyU within 10 nt of the predicted 3’ terminus (37% of terminators with predicted hairpins; Figure 1B). We used five consecutive U residues within 10 nt of the predicted 3’ terminus as a criterion for polyU, allowing for 1 alternative base at any position. PolyU length was defined as the longest continuous stretch of U residues. In contrast 15 hairpin-free termini of the termini lacking predicted hairpins include a polyU sequence (15% ), a slight enrichment of polyU located at termini with a predicted hairpin. We performed an identical analysis on the *E. coli* Term-Seq dataset, rescaling free energy from *in silico* RNA folding to a temperature of 37°C. In this case, amongst termini with predicted hairpins, 350 out of a total of 539 included a polyU within 10 nt of the predicted 3’ terminus (65% of the total termini with predicted hairpins). Amongst the candidate intrinsic terminators containing both predicted hairpin and polyU, average polyU was longer for *E. coli* (mean 5 nt) than *T. aquaticus* (mean 4 nt) (Figure 1C).

### *Taq* terminators show adaptation for thermal stability

RNA hairpin stability, and thus potentially intrinsic termination, is temperature dependent. To assess possible adaptation of *Thermus* terminators to high physiological temperature (25) we compared our predicted 3’ terminus sequences with those predicted from previously published data for the mesophile *E*. *coli* (32). We performed *in silico* RNA structure prediction with rescaling free energy for a range of temperatures from 25-90°C. As controls we used 1) dinucleotide shuffled versions of the *Taq* 3’ terminus sequences and 2) 500 randomly sampled 61 nt sequences within *Taq* CDSs, which we expected to contain virtually no functional terminators. *Taq* 3’ termini showed an apparent increased thermal stability compared with those from *E. coli* and controls (Figure 2A). The shift in predicted stability of the dinucleotide shuffled versus native *Taq* terminators was minor compared to the corresponding shift for *E. coli*.

**Figure 2.**
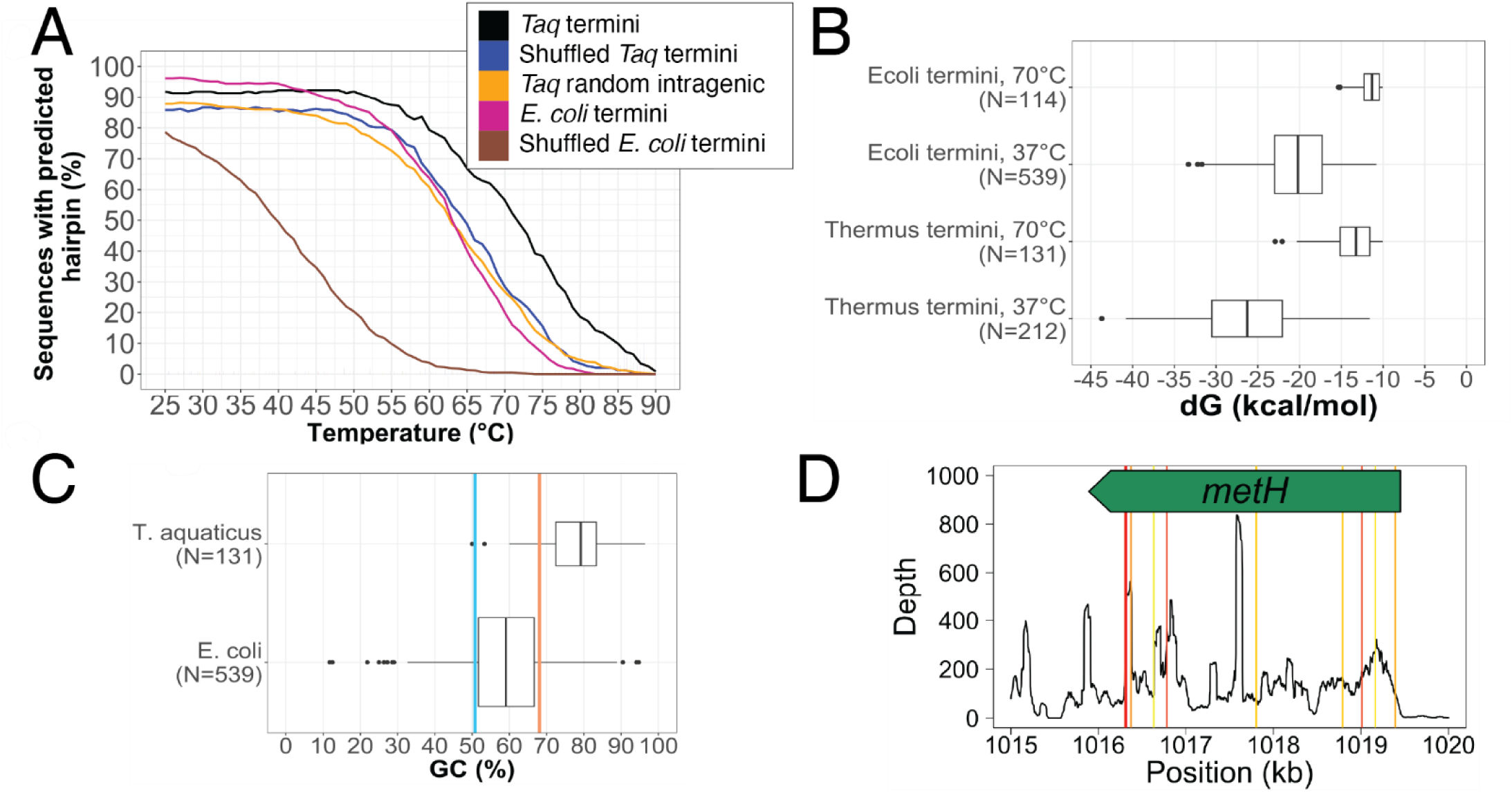
**(A-C)** *In silico* predicted hairpin stability at *E. coli* and *Taq* candidate “sharp” 3’ termini. **(A)** Proportion of 3’ termini with predicted stable hairpins when rescaling free energy to varying temperatures. “Shuffled” indicates dinucleotide shuffled version of the 3’ termini sequences. “Random” indicates 61 bp randomly selected intragenic sequences. **(B)** Predicted hairpin free energy. **(C)** GC content of predicted hairpin stem sequences for *Taq* and *E. coli* with free energy adjusted for 70°C and 37°C, respectively. Red and green lines indicated whole genome GC content for *Taq* and *E. coli*, respectively. For boxplots, box width is proportional to the square root of sample size, and sample size for each bin is presented on the x axis. (B, C) include only termini proximal to a predicted hairpin. **(D)** Example of intragenic 3’ termini, Term-Seq depth of coverage across *metH* gene shown (more examples of long genes in Fig. S4). Depth peaks indicated mRNA 3’ ends. RNA hairpins were predicted by *in silico* structure prediction of 61 bp sequence segments with a 30 bp overlap covering the length of the gene. Free energy was rescaled for RNA folding at 70°C. Predicted hairpin positions are indicated by vertical lines. Hairpins were ranked - lines colour is scaled from red to yellow, respectively indicating longer and shorter hairpins within the gene.

At physiological temperatures of *Taq* (70°C), and *E. coli* (37°C), mean predicted hairpin stem length was similar for both species; 8.8 bp for *Taq* and 10.0 for *E. col* (Figure S1). The latter is consistent with a previously reported value (33). When comparing at an equal temperature of 37°C, the mean predicted free energy amongst *Taq* termini with predicted stable hairpins was significantly lower than for *E. coli* (-26.4 kcal/mol vs -20.2 kcal/mol; p-value < 2.2×10^−16^, figure 2B).

The higher predicted stability of candidate *Taq* termination hairpins results from their enriched GC content. The mean GC content for the stem of predicted stable *Taq* terminator hairpins at 70°C () was significantly higher than that predicted for *E. coli* at 37°C (77.9% vs 59.2%; p-value < 2.2×10^−16^, Figure 2C), and was substantially higher than the overall genomic GC content of *Taq* (68.1%) (34).

### Term-Seq data analysis detected intragenic RNA 3’ ends in *Taq*

Many of the predicted *Taq* 3’ termini were located within coding sequences (263 out of 530 total), despite RNA isolated from *Taq* showed good integrity, prior to library prep for Term-Seq (Figure S2). Indeed, we noted a “noisy” aspect to the *Taq* Term-Seq data, which contained many small peaks in sequencing depth distributed across the genome. Prior to filtering for peaks with the greatest depth in each 1 kb window, signal processing identified 10396 depths peaks across the *Taq* genome (mean of three samples). In contrast, we identified only 1475 Term-Seq depth peaks in *E. coli*.

Four illustrative genes demonstrate the apparent intragenic 3’ ends - metH, dnaE and rpoC, the longest annotated CDS and a large non-coding 23S rRNA (Figure 2D and Supplementary Figure S3). The large peak downstream of the gene and other co-orientated genes was presumed to be the operon 3’ terminus. We noted a substantial number of peaks indicating potential transcript 3’ ends inside the genes. Peak heights varied, and some internal peaks exceeded those at the presumed operon end. *In silico* prediction identified RNA hairpins within genes, generally associated with Term-Seq depth peaks. However, several large Term-Seq peaks lacked predicted hairpins, and many predicted hairpins did not coincide with Term-Seq peaks.

### *Taq* RNAP terminates on native terminators *in vitro*, requiring both hairpin and polyU, but not NusG

We performed *in vitro* transcription assays with recombinant *Taq* RNAP to confirm the capacity for intrinsic termination (Figure 3). We selected two putative intrinsic terminators identified by Term-Seq, with high depth of 3’ end sequencing (>1000 reads) and highly consistent 3’ end position across reads. AP and CAT were named after their upstream genes - “aminopeptidase” (locus tag TO73_RS05025) and “manganese catalase family protein” (locus tag TO73_RS09610). *E. coli* RNAP and the well-characterised bacteriophage λ terminator tR2 (17) were tested as mesophilic controls. All terminators were placed in the same sequence context - downstream of T7A1 promoter and region coding for the 21nt long RNA (Figure 3A).

**Figure 3.**
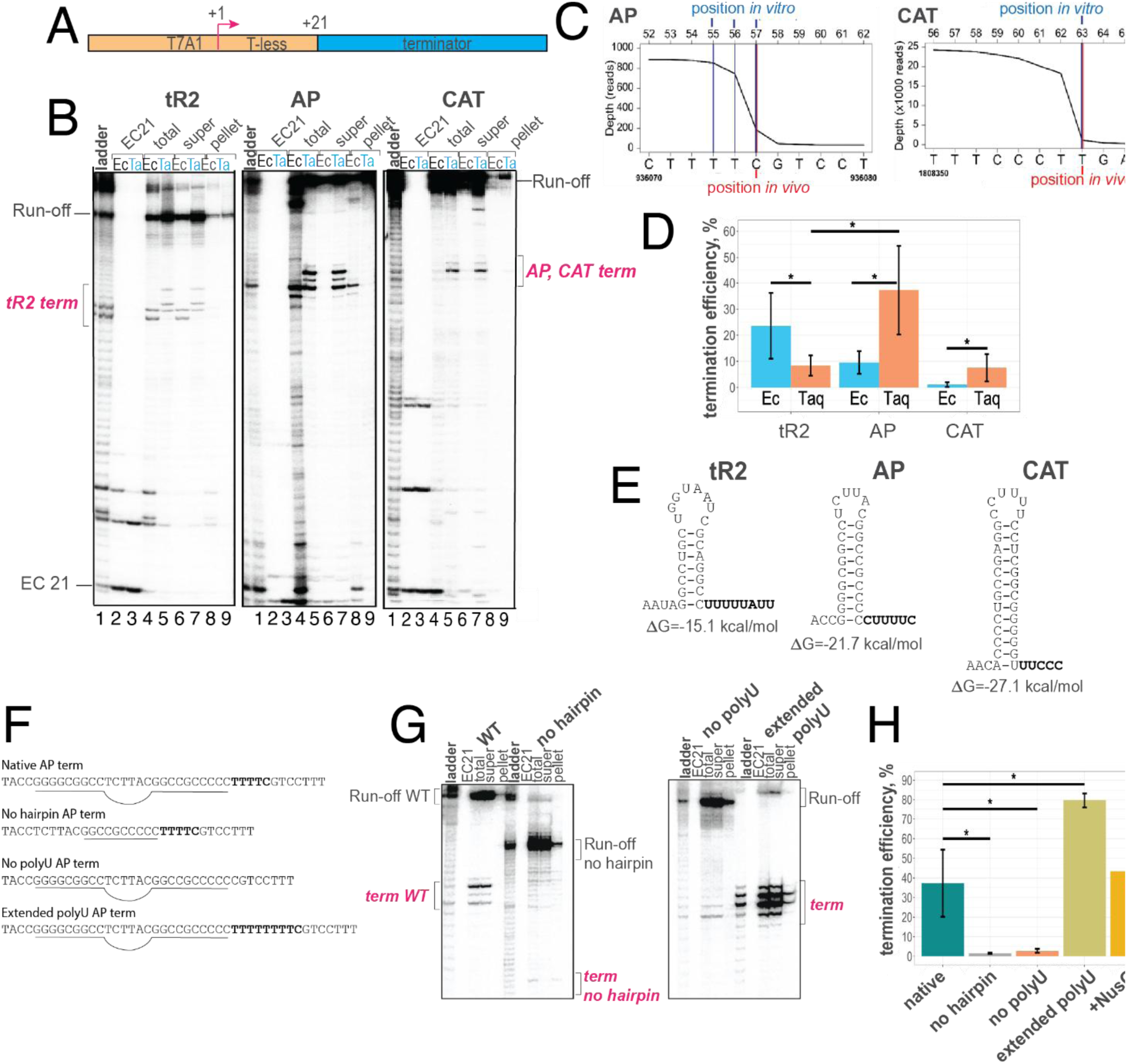
Termination at the native *Taq* AP and CAT terminators. **(A)** Scheme of the templates for *in vitro* transcription. **(B)** PAGE analysis of *in vitro* transcription products for tR2, AP and CAT terminators. Lanes: (1); template specific ladder, (2, 4, 6, 8); *E. coli* RNAP, stalled elongation complex with 21-nt long RNA, EC21, and post reaction total, supernatant and pellet fractions, respectively. (3, 5, 7, 9); *Taq* RNAP, EC 21, and post reaction total, supernatant and pellet fractions, respectively. **(C)** Term-Seq depth across the AP and CAT terminator positions. Genomic location and sequence are indicated by the bottom y-axis, and position on the *in vitro* transcription template relative to the transcription start site is indicated on the top y-axis. Termination positions estimated by Term-Seq and *in vitro* transcription are indicated by red and blue lines, respectively. **(D)** termination efficiency determined from the band intensity from PAGE analysis of *in vitro* transcription termination products. Green and red indicate *E. coli* and *Taq* RNAP, respectively. Results are presented as the mean of at least 3 independent experiments. Error bars indicate standard deviation. Asterisks indicate significant differences (p-value < 0.05, one-sided Mann-Whitney U test). **(E)** Minimum free energy structures for terminator hairpins. **(F)** Variants of AP terminator sequences, only sequences around terminator are shown. **(G)** PAGE analysis of *in vitro* transcription using templates shown on (G). **(H)** Termination efficiency on AP variants templates, the mean of at least 3 independent experiments. Error bars indicate standard deviation. Asterisks indicate significant differences, as before.

*Taq* RNAP demonstrated termination, including transcript release at all three terminators (Figure 3B). In contrast, *E. coli* was only able to terminate at the tR2 terminator and pause at AP terminator, with almost no termination for CAT. There was no detectible transcript release for *E. coli* RNAP at the AP terminator. The *Taq* RNAP termination position was shifted downstream compared to that of *E. coli* RNAP for tR2 and AP terminators. In addition, *Taq* RNAP generated three major termination products for the tr2 and AP terminators, shifted in 3’ position by 1-2 bases, compared with two for *E. coli* RNAP. For *Taq* RNAP, termination position agreed exactly between *in vivo* (Term-Seq) and *in vitro* experiments (Figure 3C). Termination efficiency for *Taq* RNAP was relatively low at given experimental conditions for the tR2 and native CAT terminators (∼10%) but was significantly higher for the native AP terminator (37%) (Figure 3D). Termination was significantly higher for *Taq* RNAP than *E. coli* RNAP for native *Taq* terminators, whereas the reverse was true for the native *E. coli* phage terminator tR2. The RNAFold predicted free energy for the minimum free energy structure of the tR2, AP and CAT terminator hairpins was -15.10 kcal/mol, -21.70 kcal/mol and -27.10 kcal/mol (Figure 3E), suggesting that the native terminator hairpins are more stable than the mesophilic tR2 terminator hairpin.

We performed additional *in vitro* experiments to further characterise factors important for *Taq* RNAP termination, using variants of the AP terminator (Figure 3F). Deletion of the upstream stem of the terminator hairpin abolished termination nearly completely (Figures 3G, 3H), demonstrating the necessity for hairpin formation. To investigate the functional importance of the polyU, we generated modified terminators lacking polyU, and with an extended 8 nt long polyU. PolyU deletion greatly reduced termination efficiency (to 4%), whereas termination was greatly enhanced for the extended polyU template (to 75%; Figures 3G, H). Transcript release efficiency was not substantially affected by polyU length. Recombinant *Taq* NusG had no significant effect on termination efficiency (Figure 3H).

### *Taq* RNAP stabilises the RNA hairpin during termination

To investigate whether temperature-dependent melting of the terminator hairpin would be a limiting factor for intrinsic termination, we measured the effect of temperature on termination efficiency for the AP terminator (Figure 4A). Unexpectedly, we observed a positive, linear relationship between temperature and termination efficiency (Spearman correlation coefficient 0.96) in the range of 40-80°C (the physiological growth range of *Taq* (25)). Above 80°C RNAP lost its activity. 40-80°C temperature variation resulted in a buffer pH decrease from 7.8 to 7.1. No significant difference was detected between pH 7.0 and 7.8 at 66.5°C, suggesting that the major effect on termination efficiency resulted from temperature alone (Supplementary Figure S4). We hypothesized that low temperatures may increase R-loop formation behind elongating RNAP, disrupting terminator hairpin folding and termination. To detect R-loops, we used an RNase H, which selectively degrades RNA within RNA-DNA hybrids. (Supplementary Figure S5) Minor RNase H degradation products were detected at 40°C, but not 66.5°C, suggesting R-loop formation in a small proportion of products. However, the run-off transcription products were unaffected by RNase H, suggesting that the majority of unterminated transcripts did not form R-loops.

**Figure 4.**
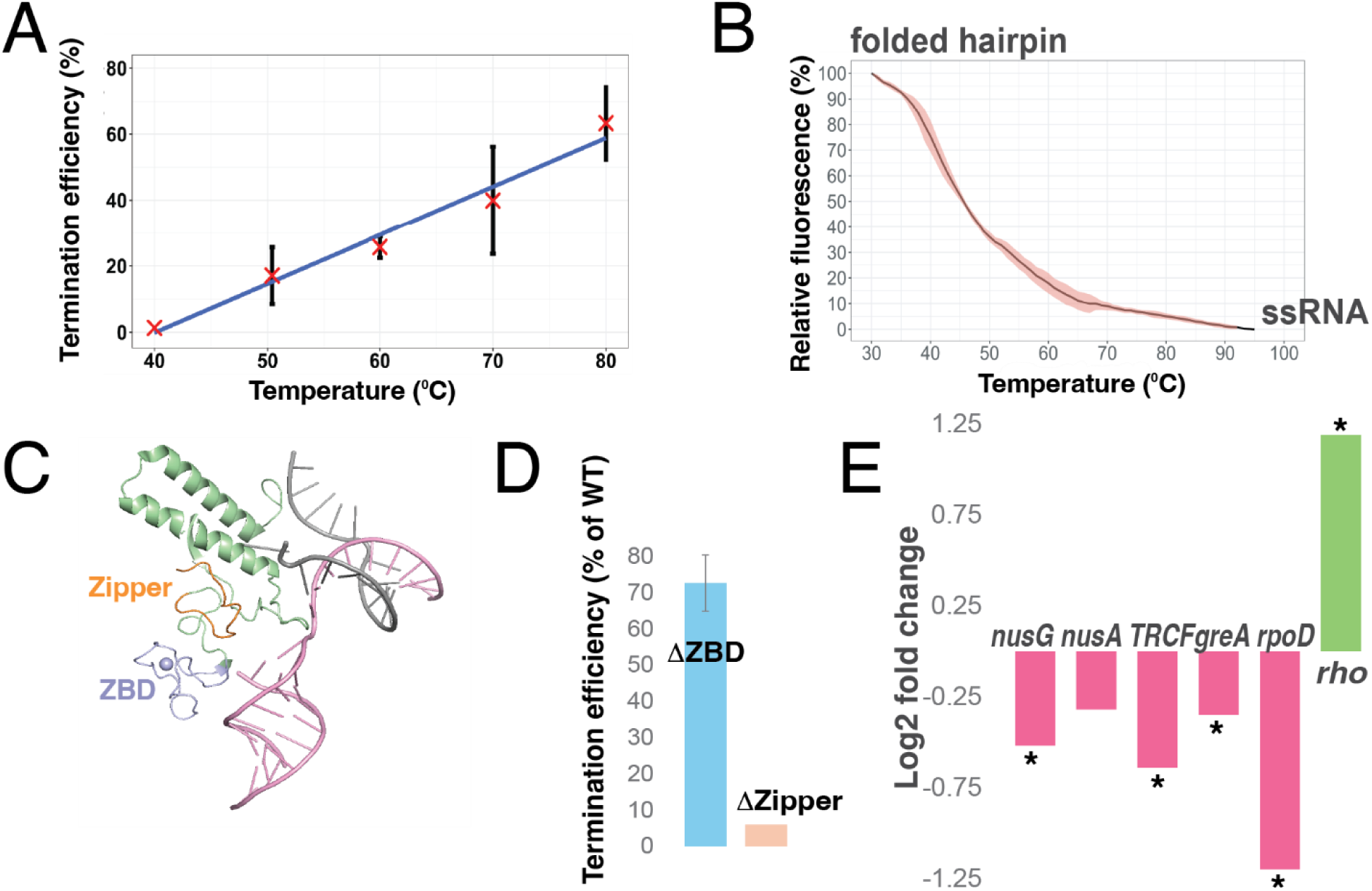
*Taq* RNAP stabilises the terminator RNA hairpin. **(A)** Termination efficiency (determined from the band intensity from PAGE analysis of *in vitro* transcription termination products) of *Taq* RNAP at the native AP terminator with varying temperature. Results are presented as the mean of 3 independent experiments. Error bars indicate standard deviation. The fitted linear regression model is presented in blue. **(B)** Melting analysis of AP terminator hairpin ssRNA oligonucleotide, determined by EvaGreen® binding assay. Results are presented as the mean of 3 independent experiments. Red shaded region indicates standard deviation. **(C)** Structures of the three protein loops surrounding RNA duplex in hairpin complex in *E. coli* pre-termination elongation complex, adapted from PDB file 7YPA. **(D)** Deletions of zink-binding domain (ZBD) and Zipper domain affect termination efficiency by *Taq* RNAP, plotted in % from efficiency of WT RNAP (determined from the band intensity from PAGE analysis of *in vitro* transcription termination products as before). For ΔZBD RNAP the mean is shown of 3 independent experiments. Error bars indicate standard deviation. For ΔZipper RNAP the experiment was done twice. **(E)** Differential expression of transcription factors genes of *Taq* grown at 55 °C vs *Taq* grown at 70 °C, log_2_ fold change. The results are the mean from three independent biological replicates in each condition. Genes identified as significantly differentially expressed are indicated with an asterisk (false discovery rate < 0.05).

To determine the intrinsic thermal stability of the AP terminator hairpin independent of the transcription elongation complex, we measured its melting behavior using high-resolution thermal denaturation analysis with the double-stranded RNA–binding dye EvaGreen (35) under buffer conditions of *in vitro* transcription (Figure 4B). We used the synthetic ssRNA oligonucleotide corresponding for the hairpin-forming and polyU parts of the AP terminator. The results suggested that on its own the hairpin is almost entirely unfolded at 80 °C, the temperature at which *Taq* RNAP showed maximal termination efficiency.

These results suggest that RNAP facilitates terminator hairpin folding. Formation of the terminator hairpin occurs within an RNA exit channel formed by protein loop domains, including the Zipper and the zinc-binding domain (ZBD) (Figure 4C). Deletion of Zipper domain resulted in near-complete loss of termination on AP terminator, whereas deletion of the ZBD decreased termination by approximately 25% (Figure 4D).

### Expression of Rho termination factor is increased with decreasing temperature

How Taq RNAP achieves transcription termination at lower temperatures within the physiological range of *Taq*, where intrinsic termination is predicted to be increasingly inefficient, is unknown. One possibility is that Rho-dependent termination assumes a larger role under these conditions. We analysed differential gene expression in *Taq* grown at 55 °C vs 65 °C. Principal coordinate analysis suggested a high degree of reproducibility between samples, with a distinct transcriptional profile between the two temperatures (Supplementary Figure S6). At 55 °C, cells showed reduced expression of key transcription factors genes, including the initiation factor sigma (*rpoD*), the elongation factors *nusA*, *nusG*, *greA* and the repair factor *TRCF*, the Mfd homolog. In contrast, expression of the termination factor Rho significantly increased (Figure 4E). This expression pattern is consistent with an increased contribution of Rho-dependent termination under lower-temperature conditions. Attempts to directly test Rho dependence using the inhibitor bicyclomycin were unsuccessful due to rapid compound inactivation (within 20 min) at 55 °C.

## DISCUSSION

This work represents genome-wide analysis of transcript 3’ termini in a thermophilic bacterium. Our data demonstrate that the canonical intrinsic termination mechanism based on a nascent RNA hairpin and a weak U-rich RNA–DNA hybrid functions *in vivo* and *in vitro* at ∼70 °C in *Taq*.

We found that intrinsic termination is a major mechanism for *Taq*, as RNA hairpins are predicted near approximately half of 3’ termini, similar to *E. coli* (11). The remaining fraction of 3’ termini may represent Rho-dependent termination and non-hairpin pause elements (36). The presence of termini with highly variable 3’-end location across the Term-Seq reads, termed “soft” terminators, likely results from transcript 3’-ends processing (37).

Our Term-Seq analysis revealed that *Taq* had a lower proportion of termini with proximal polyU sequence, and that where present the polyUs were on average shorter than mesophile *E. coli*. Similar *in silico* predictions have been made for potential intrinsic terminators in other bacteria with high GC content genomes (38). Possibly the shortened polyUs found amongst native *Taq* terminators results from a trade-off between termination efficiency, and the selection pressure which had promoted a high-GC genome. The role of a long polyU could be less significant at higher temperatures, as the DNA-RNA hybrid is likely to more readily dissociate. On the other hand, we might have underestimated the number of polyUs due to the parameters we used for polyU identification during Term-Seq data analysis. Still, the results from our *in vitro* experiments support the hypothesis that the polyU is functionally important for *Taq* termination, as deletion of the polyU greatly reduced termination efficiency and its extension – increased, and native CAT terminator with shorter polyU was less efficient despite having stronger hairpin.

A central question concerns the behaviour of the termination hairpin at elevated temperature. Our analysis indicates that hairpins proximal to *Taq* terminators are optimized for enhanced thermal stability, largely via enriched GC content, similarly to what has been shown for rRNA and tRNA structures of thermophilic bacteria (24). The enhanced *in vitro* termination of *Taq* RNAP at the native AP terminator compared with the mesophilic tR2 terminator could suggest that greater hairpin stability plays a beneficial role in termination. Also, the independent contribution of sequence composition alone suggested by *in vitro* results for *E. coli* (39) and *T. thermophilus* (26), was also demonstrated here by decrease in predicted stability of the dinucleotide-shuffled terminator hairpins. It is unclear whether the observed thermal stability adaptations of the terminator hairpins may function to protect released transcript against degradation (40).

Increased hairpin stem length have been hypothesised as an alternative mechanism of thermal adaptation (41). However, our results did not show any difference in average hairpin stem length between *Taq* and the mesophile *E. coli*. We suggest that requirement to accommodate hairpin in the RNA exit channel places strong constraints on functional hairpin stem length range, reminiscent of the proposed allosteric model of intrinsic termination (42).

Somewhat surprisingly, our *in vitro* data did not demonstrate that high temperature is a limiting factor for intrinsic termination. We showed a positive linear relationship between termination efficiency and temperature. A similar trend has been reported for *E. coli* RNAP (43), however, in contrast to *E. coli*, the temperature range examined here is physiologically relevant for the thermophilic *Taq*. We suggest that in general, a higher temperature will promote the entropically-driven termination pathway (44) through disassociation of the molecular components of elongation complex.

Moreover, our results showed that the temperature for optimal termination efficiency was substantially above the temperature at which the free RNA oligonucleotide containing terminator hairpin was entirely melted in solution. We take it as the direct experimental evidence that RNAP provides RNA chaperone activity to stabilise the terminator hairpin. This function was also hypothesised for the zinc finger, zipper and flap domains of RNAP in *E. coli* RNAP (15,42,45). Indeed, we demonstrated that deletion of the ZBD and especially Zipper domain of *Taq* RNAP negatively affected termination efficiency.

We propose that Rho-dependent termination plays an enhanced role near the lower bound of *Taq*’s physiological temperature range, where intrinsic terminator efficiency is reduced. Intrinsic termination can, in some cases, be assisted by Rho factor (46). Accordingly, some terminators identified here may operate through this “mixed” mechanism. Direct analysis of Rho-dependent termination in *Taq* has proven difficult due to thermal instability of the Rho inhibitor bicyclomycin. Predictive tool RhoTermPredict (46) seem to overestimate Rho-dependent terminators’ ubiquity (including within coding regions), likely due to genome composition. Nevertheless, our RNA-Seq data demonstrated that at 55 °C expression of Rho factor is significantly elevated. This suggests that Rho translocase activity may become increasingly important under these conditions, supported by: (i) Rho expression is specifically higher at lower temperatures, when overall promoter activity declines due to less efficient promoter DNA melting and lower level of primary initiation factor expression, and (ii) reduced expression of elongation factors at 55 °C suggests a lower abundance of elongating RNAP complexes, increasing the ratio of Rho to transcription complexes.

We found no enhancement of *Taq* RNAP termination efficiency by NusG, in agreement with previous results for *Thermus thermophilus* RNAP *in vitro* (27).

In summary, our results provide an atlas for transcript termini across the *Taq* genome, and suggest intrinsic termination is a major termination mechanism in *Taq*. Transcript termini hairpins appear to be optimised for thermal stability, but we did not find any evidence that high temperature is a limiting factor for termination via hairpin instability. Together, our evidence strongly suggests an RNA chaperone activity for RNAP and temperature-dependent shift between alternative termination mechanisms.

## MATERIALS AND METHODS

### Strains and culture conditions

For Term-Seq and RNASeq analysis, overnight cultures of *Thermus aquaticus* strain YT-1 were diluted to an OD_600_ of 0.1 and, incubated at 70°C (unless otherwise stated), with shaking at 180 rpm. Cultures were harvested in 10 ml aliquots between OD_600_ 0.5-0.7 by centrifugation at 1500 x g for 15 minutes at 4°C, immediately suspended in 750 µl RNALater (Thermo Fisher Scientific), flash frozen in liquid nitrogen and stored at -70°C for no longer than 1 month before RNA extraction.

### RNA isolation

Samples were diluted 1:2 with PBS and cells pelleted by centrifugation at 1500 × g for 15 min at 4°C. Pellets were resuspended in RNAPro (MP Biomedicals) and lysed by bead beating (6500 rpm, 3 × 30 s, 30 s breaks, 4°C) using lysing matrix B tubes. Lysates were cleared at 12,000 × g for 5 min at 4°C and incubated 5 min at room temperature. Chloroform (300 µl) was added, vortexed, and incubated 5 min before centrifugation at 12,000 × g for 5 min at 4°C. The aqueous phase was collected, and RNA was purified using the Norgen Total RNA Purification Plus kit.

### Term-Seq

Term-Seq library preparation and sequencing were performed by Vertis Biotechnologie AG. RNA integrity was checked using a MultiNA microchip electrophoresis system (Shimadzu). Total RNA was subjected to rRNA-depletion by an in-house method, followed by ligation of 5’ Illumina sequencing adapters to the RNA 3’ ends. M-MLV reverse transcriptase was used to perform first-strand cDNA synthesis, followed by fragmentation and ligation of the 3’ Illumina sequencing adapter 3’ cDNA end. cDNA fragments were amplified by PCR using a high-fidelity polymerase, and products were purified using the Agencourt AMPure XP kit (Beckman Coulter Genomics). Barcoded cDNA samples were pooled and sequenced on a NextSeq 500 instrument (Illumina) to generate 10-12 million single-end 75 bp reads per sample. For comparison, the published dataset for *E. coli* Term-Seq (32) was downloaded from the SRA database (47) under accession SRR6519353-SRR6519355.

### RNA-Seq

RNA-Seq library preparation and sequencing were performed by Novogene (Prokaryotic RNA-Seq service). RNA sample integrity was assessed using the Agilent 5400 fragment analyser system. For library preparation, first rRNA was depleted, followed by fragmentation, reverse transcription, second strand cDNA synthesis, end-repair and A-tailing, adapter ligation, USER enzyme digestion, fragment size selection and PCR amplification. Resulting libraries were sequenced using a Illumina Novoseq X plus instrument to generate 150 bp paired end reads.

### Identification of transcript 3’ termini

Low-quality reads with >0.25 bp expected errors were removed using Usearch v11.0.667 (48), and Illumina adapters were trimmed with Cutadapt v4.2 (49). Because the available *Thermus aquaticus* YT-1 genome was highly fragmented, reads were aligned instead to the contiguous *T. aquaticus* Y51MC23 reference genome (NCBI ASM139977v1) (34) using Bowtie2 v2.4.2 (50). *E. coli* Term-Seq datasets (SRA accessions SRR6519353–SRR6519355) (47) were aligned to the BW25113 reference genome (GenBank CP009273.1) (51) using the same pipeline.

Samtools v1.10 (52) was used for alignment file manipulation and depth-of-coverage calculations. Custom R scripts identified transcription terminators. Peaks in Term-Seq coverage, representing RNA 3′ termini, were detected using the peaks function from the IDPmisc R package. For *T. aquaticus*, the depth threshold for peak detection was 35 reads, and minimum peak height was 50 reads. Because sequencing depth for the *T. aquaticus* dataset was estimated to be ∼3.6-fold higher than that of *E. coli*, thresholds for *E. coli* were reduced by dividing by 5.6 to maintain comparable sensitivity.

To identify the most abundant 3′ end for each transcript, only the peak with the highest depth within a 1-kb window was retained. Peaks were classified as “sharp” (sharp transitions, fixed 3′ ends) or “soft” (variable ends, plateau or parabolic shapes). Within 75 bp of each peak, if the change in coverage over 1 nt exceeded 50% of the maximum peak height, the terminus was labeled “sharp.” For defined peaks, the 3′ end was assigned to the position with the greatest coverage drop within 150 nt downstream. For soft peaks, the 3′ end was assigned 38 nt downstream (half the read length).

Termini had to be detected within 20 nt across all three biological replicates. Terminus position and termination efficiency were averaged between replicates, and terminus type (“sharp” or “soft”) was assigned by majority rule.

Sequences spanning 50 nt upstream and 10 nt downstream of each terminus were analyzed for RNA secondary structure using RNAfold v2.5.1 (53). A stable hairpin required a ≥4-nt stem and predicted free energy < –10 kcal/mol. Poly-U tracts were defined as ≥5 consecutive U residues within 10 nt of the 3′ end, allowing one non-U substitution; poly-U length was the longest continuous U stretch. Statistical significance was assessed via the Mann-Whitney U test.

Statistical significance (p-values) of the mean predicted free energy and hairpins length differences between *E. coli* and *T. aquaticus* were calculated using two-tailed Mann-Whitney U test.

### Analysis of Differential Gene Expression

Differential gene expression was assessed from RNASeq data. Reads were aligned to the *Thermus aquaticus* strain Y51MC23 reference genome (NCBI assembly accession ASM139977v1) (34) using Bowtie2 version 2.4.2 (50). Samtools version 1.10 (52) was used to manipulate alignment files and calculate depth of coverage across the reference genome. Aligned reads mapping to annotated features were counted using HTSeq version 0.9.1 with default settings. Differentially expressed genes were identified via the edgeR version 3.42.4 (54) pipeline, using default settings for filtering low-expressed genes, normalising for library size, estimating negative binomial dispersions and performing the exact test for differential expression.

### Protein purification

Recombinant *Taq* RNAP and σ^A^ were isolated as described (55). *Taq nusG* gene was cloned into pET28a vector and expressed in T7express strain (NEB). Cells pellets were resuspended in lysis buffer (50 mM Tris HCl pH 8.0, 250 mM NaCl, 10% glycerol, 5 mM imidazole, and protease inhibitors cocktail (Roche). Cells were lysed by sonication, lysate clarified by centrifugation for 15 minutes at 15 kg. Lysate was incubated at 70°C for 30 minutes, soluble fraction was collected after centrifugation for 15 minutes at 15 kg and applied to 5 ml HisTrap column (Cytiva). Fractions containing NusG were eluted with lysis buffer containing 200 mM imidazole, analysed by SDS PAGE, pooled, concentrated using Amicon Ultra 3 kDa cut-off centrifugal device, and dialysed overnight against storage buffer (40 mM Tris HCl pH 8.0, 200 mM KCl, 50% glycerol, 1 mM EDTA, 1 mM DTT).

### *In vitro* transcription

*In vitro* transcription was performed on dsDNA templates containing T7A1 promoter upstream of 21 nucleotides common sequence, followed by the specific terminator sequences (Fig. 3A and Supplementary file 1).

*Taq* RNAP open promoter complexes were prepared by incubating 235 nM RNAP, 165 nM biotinylated DNA and 7 µM *Taq* σ^A^ in initiation buffer (40 mM TrisHCl, 80 mM KCl, 20 mM MgCl_2_, pH 7.7 at 66.5°C) for 10 minutes. Buffer. Elongation complexes containing radiolabelled 21-nt transcript were prepared in with 87 nM open promoter complex in transcription buffer (40 mM KCl, 10 mM MgCl_2_, 20 mM TrisHCl, pH 8.00), 20 µM ATP and CTP, 11 µM CAUC and 0.4 µM [α-^32^P] GTP (4000Ci/mmol). Elongation was allowed to proceed at 66.5°C for 3 minutes. Reactions were cooled to RT for 1 minutes, complexes were immobilised on 7% [v/v] streptavidin Sepharose (Cytiva) at RT for 2 minutes. 1.1 µM cold GTP was added for 15 s. Immobilised complexes were washed twice with 1 ml of buffer containing 1 M KCl, 10 mM MgCl_2_, 20 mM TrisHCl pH 8.00, twice with transcription buffer, and once with “physiological” buffer (100 mM potassium glutamate, 33 mM TrisHCl, pH 7.6 at 66.5°C).

Reactions (25 µl) after prewarming for 2 minutes were supplemented with 250 µM NTPs and allowed to proceed for 2 minutes. Reaction fractions were combined with an equal volume of stop buffer (7M urea, 0.02% bromophenol blue, 0.02% xylene cyanol, 22 mM EDTA, 100 µg/ml heparin, 46% formamide, 89 mM Tris-borate). “Supernatant” fractions were prepared after pelleting the resin, “pellet” fraction - by washing the pellet twice with 1 ml transcription buffer.

*E. coli* RNAP assays were performed identically, except that 700 nM recombinant *E. coli* σ^70^ was used at 37°C (the pH of initiation and physiological buffers was adjusted to 7.7 and 7.6 at 37°C).

Template-specific RNA ladders with 1 nucleotide resolution were prepared by hydrolysing “total” fractions in 0.67X alkaline hydrolysis buffer (Ambion) at 95°C for 1 minute 45 seconds, followed by neutralisation with an equal volume of stop buffer.

Transcription products were resolved by denaturing PAGE, revealed by PhosphorImaging (Cytiva) and analysed using ImageQuantTL v10.2, Cytiva. Termination efficiency was calculated as the percentage intensity of the termination band(s) as a fraction of all bands beyond (and including) the termination band(s).

RNase H analysis of R-loop formation was performed in 7.5 µl reaction volumes containing 100 units/ml thermostable RNase H (NEB). Positive control reactions included RNase H and 840 nM ssDNA oligonucleotide complimentary to the 3’ end of the run-off transcript (sequence 5’-GTCATCTGTAGCGTGCGATCGCCCC-AAAGGAC-3’).

### RNA melting analysis

The native *vitro*AP terminator hairpin-forming RNA oligonucleotide (sequence 5’-UACCGGGGCGGCCUCUUACGGCCGCCCCCUUU-3’) were pre-folded by incubating at 80°C for 10 minutes, followed by cooling on ice for 20 minutes. Hairpin melting temperature was assessed in 10 µl reactions containing 1 µM RNA oligonucleotide and 2 µM EvaGreen® (Jena Bioscience) in physiological transcription buffer. Melting analysis was performed in a Rotor-Gene Q qPCR instrument by heating from 30°C to 95°C at a rate of 1°C every 5 seconds. Fluorescence intensity values were normalised by subtracting the signal measured at 95°C, before expressing as a percentage of the normalised signal at 30°C.

## Acknowledgments

This work was supported by Biotechnology and Biological Sciences Research Council [BB/W017385/1] to YY. Funding for open access charge: Biotechnology and Biological Sciences Research Council.

## Data Availability

The Term-Seq and RNA-Seq data is available from the SRA database (47) under accessions SRR34520674-SRR34520676 and SRR35751311-SRR35751313, respectively.

## Author Contributions

NB: Conceptualisation, Investigation, Formal analysis, Methodology, Visualisation, Writing – original draft and Writing – review & editing. ARB: Investigation. YY: Conceptualisation, Funding acquisition, Project administration, Supervision and Writing – review & editing.

## Supplementary figures

**Supplementary figure 1.**
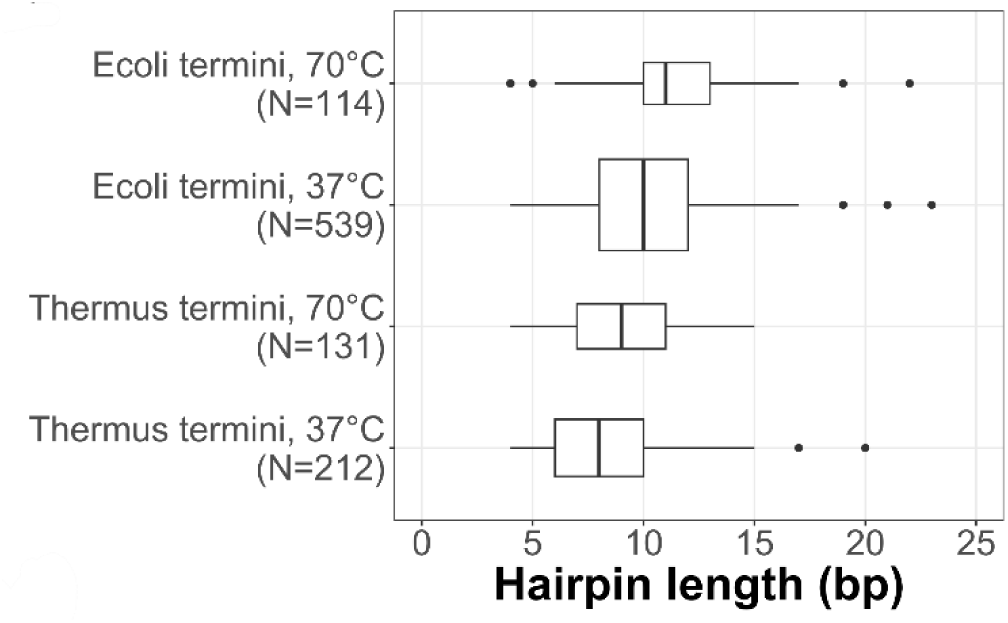
*In silico* predicted hairpin stability at *E. coli* and *Taq* candidate “sharp” 3’ termini. Hairpin length.

**Supplementary Figure 2.**
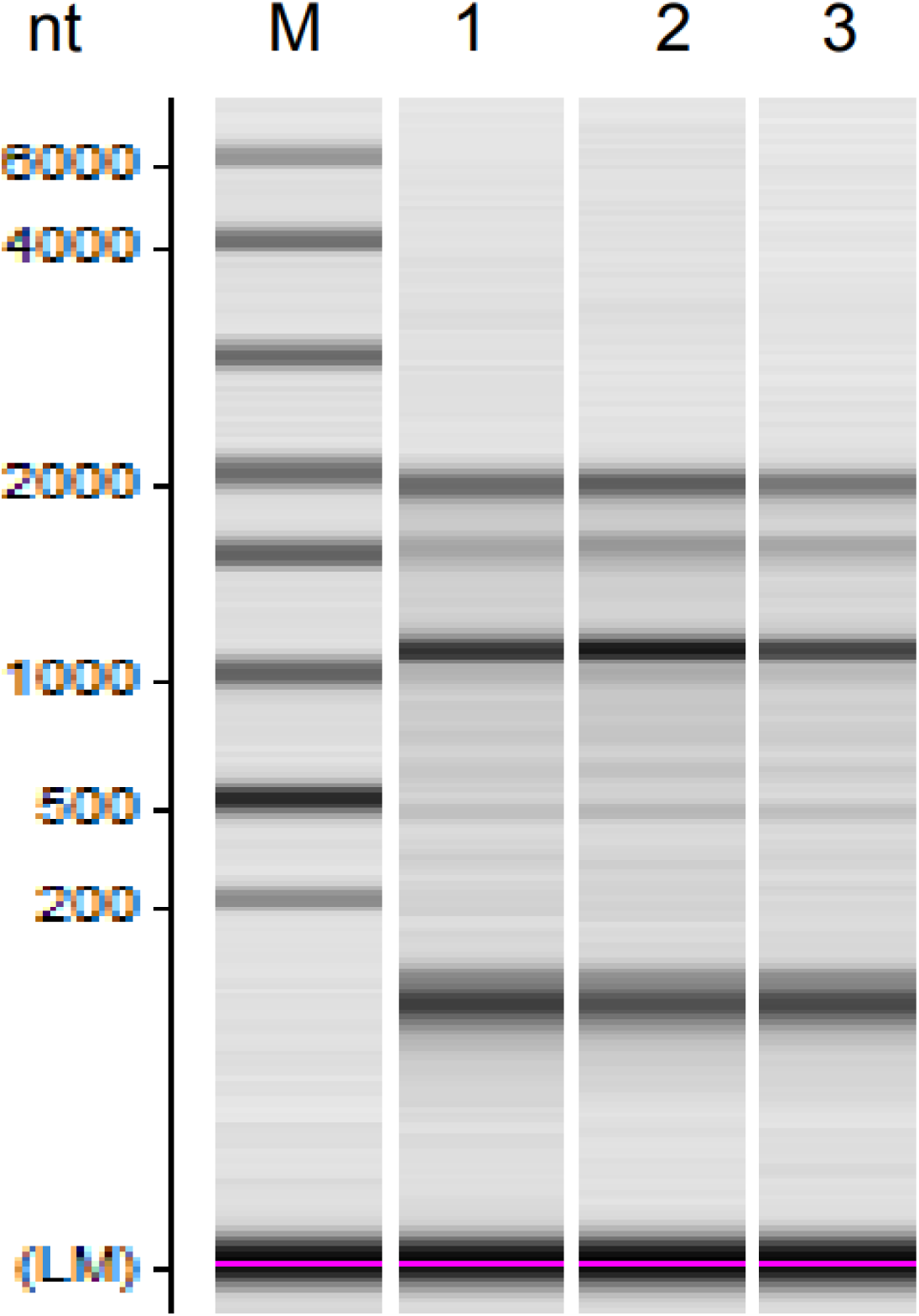
Integrity of total RNA isolated from cultured *Taq* cells assessed using a MultiNA microchip electrophoresis system (Shimadzu). M indicates size RNA markers and lanes 1-3 represent 3 biological replicates.

**Supplementary figure 3.**
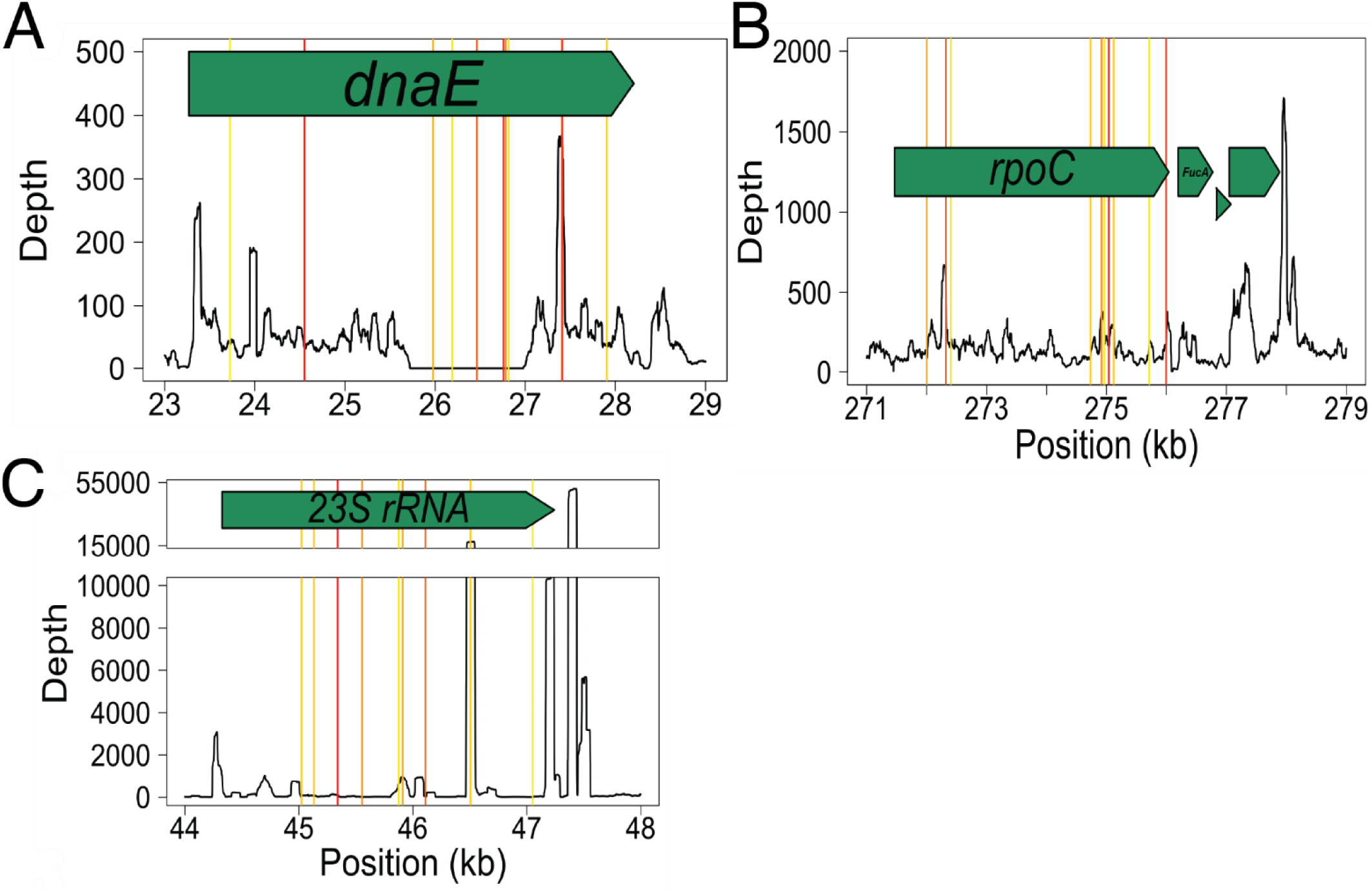
(B) Native AP terminator sequence, with the deleted upstream hairpin stem region and deleted polyU tract coloured in red and turquoise, respectively. Modified AP terminator with extended polyU is shown below. Approximate termination positions determined by *in vitro* transcription for upstream hairpin deletion, polyU deletion and extended polyU templates are highlighted in red, turquoise and grey, respectively. (C) Predicted minimum free-energy structure of the upstream hairpin deletion AP transcript coloured according to base pairing probabilities. The 3’ end corresponds to the weak *in vitro* termination product.

**Supplementary figure 4.**
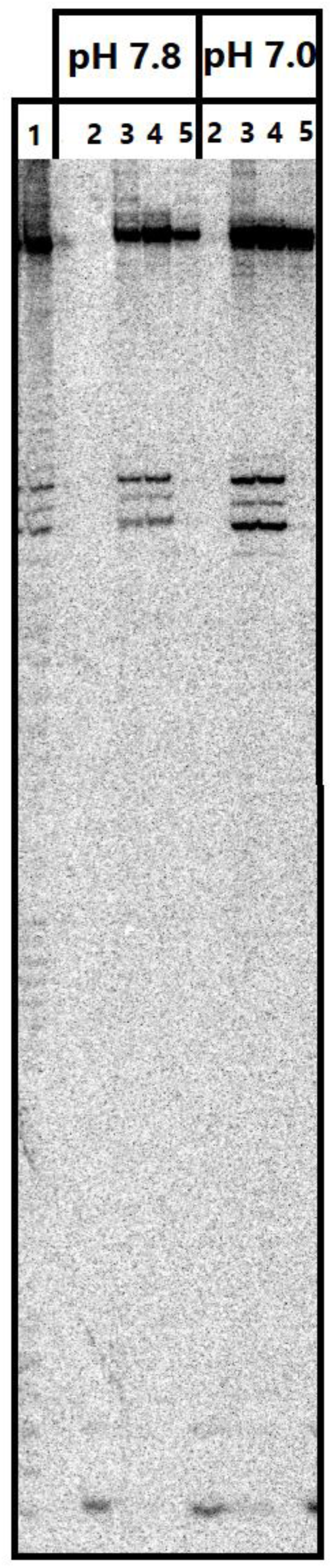
Termination efficiency at varying pH. PAGE analysis of *in vitro* transcription assay for *Taq* RNAP at the native AP terminator. Assay was performed at 66.5°C with varying pH. Lanes: (1, 2, 3, 4, 5); template specific ladder, pre-assay 21-mer control, and post assay total, supernatant and pellet fractions, respectively. pH 7.0 (mean 31%; standard deviation 3%) and 7.8 (mean 24%; standard deviation 4%)

**Supplementary figure 5.**
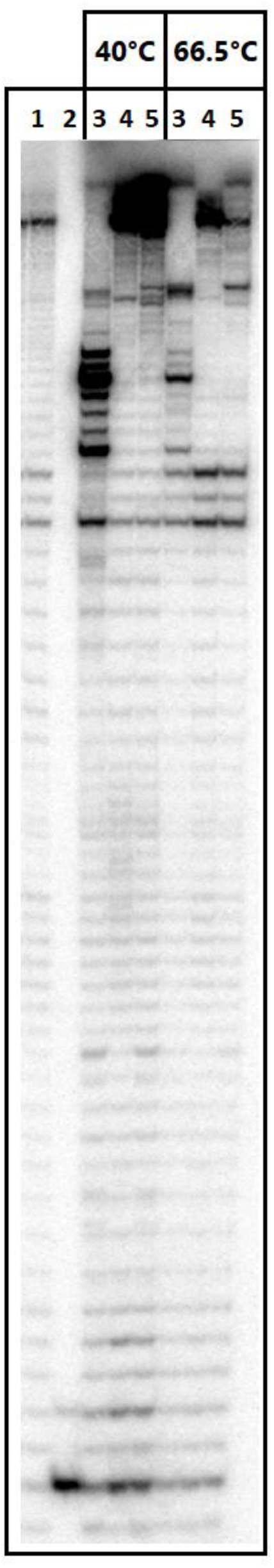
Analysis of r-loop formation. PAGE analysis of *in vitro* termination assay for *Taq* RNAP at the native AP terminator. Transcription was performed in the presence (lane 5), and absence (lane 4) of thermostable rnase H, which selectively degrades the DNA-RNA duplex of r-loops. Positive control reactions for rnase H activity (lane 3) contained an ssDNA oligonucleotide complimentary to the 3’ end of the run-off transcript. Lanes 1 and 2 were loaded with template-specific ladder and pre-assay 21-mer control, respectively.

**Supplementary figure 6.**
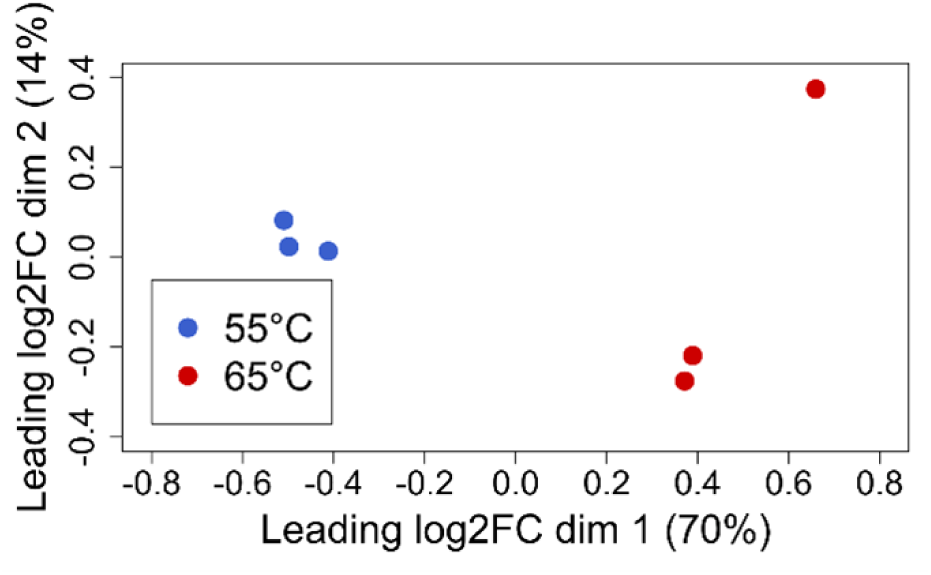
Principal coordinate analysis showing the pairwise leading log_2_ fold changes for the top 500 most differentially expressed genes between samples. The percentage of variation explained by each dimension is indicating in the axis label.

## REFERENCES

1. Platt, T. (1981) Termination of transcription and its regulation in the tryptophan operon of E. coli. Cell, 24, 10–23.

2. Yarnell, W.S. and Roberts, J.W. (1999) Mechanism of intrinsic transcription termination and antitermination. Science, 284, 611–615.

3. Lau, L.F., Roberts, J.W. and Wu, R. (1983) RNA polymerase pausing and transcript release at the lambda tR1 terminator in vitro. J Biol Chem, 258, 9391–9397.

4. Epshtein, V., Dutta, D., Wade, J. and Nudler, E. (2010) An allosteric mechanism of Rho-dependent transcription termination. Nature, 463, 245–249.

5. Molodtsov, V., Wang, C., Firlar, E., Kaelber, J.T. and Ebright, R.H. (2023) Structural basis of Rho-dependent transcription termination. Nature, 614, 367–374.

6. Yakhnin, A.V. and Babitzke, P. (2002) NusA-stimulated RNA polymerase pausing and termination participates in the Bacillus subtilis trp operon attenuation mechanism invitro. Proc Natl Acad Sci U S A, 99, 11067–11072.

7. Yakhnin, A.V. and Babitzke, P. (2010) Mechanism of NusG-stimulated pausing, hairpin-dependent pause site selection and intrinsic termination at overlapping pause and termination sites in the Bacillus subtilis trp leader. Mol Microbiol, 76, 690–705.

8. Ha, K.S., Toulokhonov, I., Vassylyev, D.G. and Landick, R. (2010) The NusA N-terminal domain is necessary and sufficient for enhancement of transcriptional pausing via interaction with the RNA exit channel of RNA polymerase. J Mol Biol, 401, 708–725.

9. Brennan, C.A., Dombroski, A.J. and Platt, T. (1987) Transcription termination factor rho is an RNA-DNA helicase. Cell, 48, 945–952.

10. Ciampi, M.S. (2006) Rho-dependent terminators and transcription termination. Microbiology (Reading), 152, 2515–2528.

11. Ahmad, E., Mahapatra, V., Vanishree, V.M. and Nagaraja, V. (2022) Intrinsic and Rho-dependent termination cooperate for efficient transcription termination at 3’ untranslated regions. Biochem Biophys Res Commun, 628, 123–132.

12. McDowell, J.C., Roberts, J.W., Jin, D.J. and Gross, C. (1994) Determination of intrinsic transcription termination efficiency by RNA polymerase elongation rate. Science, 266, 822–825.

13. Chan, C.L., Wang, D. and Landick, R. (1997) Multiple interactions stabilize a single paused transcription intermediate in which hairpin to 3’ end spacing distinguishes pause and termination pathways. J Mol Biol, 268, 54–68.

14. Yin, H., Artsimovitch, I., Landick, R. and Gelles, J. (1999) Nonequilibrium mechanism of transcription termination from observations of single RNA polymerase molecules. Proc Natl Acad Sci U S A, 96, 13124–13129.

15. You, L., Omollo, E.O., Yu, C., Mooney, R.A., Shi, J., Shen, L., Wu, X., Wen, A., He, D., Zeng, Y. et al. (2023) Structural basis for intrinsic transcription termination. Nature, 613, 783–789.

16. Gusarov, I. and Nudler, E. (1999) The mechanism of intrinsic transcription termination. Mol Cell, 3, 495–504.

17. Komissarova, N., Becker, J., Solter, S., Kireeva, M. and Kashlev, M. (2002) Shortening of RNA:DNA hybrid in the elongation complex of RNA polymerase is a prerequisite for transcription termination. Mol Cell, 10, 1151–1162.

18. Roberts, J.W. (2019) Mechanisms of Bacterial Transcription Termination. J Mol Biol, 431, 4030–4039.

19. Santangelo, T.J. and Roberts, J.W. (2004) Forward translocation is the natural pathway of RNA release at an intrinsic terminator. Mol Cell, 14, 117–126.

20. Larson, M.H., Greenleaf, W.J., Landick, R. and Block, S.M. (2008) Applied force reveals mechanistic and energetic details of transcription termination. Cell, 132, 971–982.

21. Gusarov, I. and Nudler, E. (1999) The mechanism of intrinsic transcription termination. Mol Cell, 3, 495–504.

22. Santangelo, T.J., Cubonova, L., Skinner, K.M. and Reeve, J.N. (2009) Archaeal intrinsic transcription termination in vivo. J Bacteriol, 191, 7102–7108.

23. Sanders, T.J., Wenck, B.R., Selan, J.N., Barker, M.P., Trimmer, S.A., Walker, J.E. and Santangelo, T.J. (2020) FttA is a CPSF73 homologue that terminates transcription in Archaea. Nat Microbiol, 5, 545–553.

24. Jegousse, C., Yang, Y., Zhan, J., Wang, J. and Zhou, Y. (2017) Structural signatures of thermal adaptation of bacterial ribosomal RNA, transfer RNA, and messenger RNA. PLoS One, 12, e0184722.

25. Brock, T.D. and Freeze, H. (1969) Thermus aquaticus gen. n. and sp. n., a nonsporulating extreme thermophile. J Bacteriol, 98, 289–297.

26. Berdygulova, Z., Esyunina, D., Miropolskaya, N., Mukhamedyarov, D., Kuznedelov, K., Nickels, B.E., Severinov, K., Kulbachinskiy, A. and Minakhin, L. (2012) A novel phage-encoded transcription antiterminator acts by suppressing bacterial RNA polymerase pausing. Nucleic Acids Res.

27. Sevostyanova, A. and Artsimovitch, I. (2010) Functional analysis of Thermus thermophilus transcription factor NusG. Nucleic Acids Res, 38, 7432–7445.

28. Bennett, B.D., Kimball, E.H., Gao, M., Osterhout, R., Van Dien, S.J. and Rabinowitz, J.D. (2009) Absolute metabolite concentrations and implied enzyme active site occupancy in Escherichia coli. Nat Chem Biol, 5, 593–599.

29. Berdygulova, Z., Esyunina, D., Miropolskaya, N., Mukhamedyarov, D., Kuznedelov, K., Nickels, B.E., Severinov, K., Kulbachinskiy, A. and Minakhin, L. (2012) A novel phage-encoded transcription antiterminator acts by suppressing bacterial RNA polymerase pausing. Nucleic Acids Res, 40, 4052–4063.

30. Mondal, S., Yakhnin, A.V., Sebastian, A., Albert, I. and Babitzke, P. (2016) NusA-dependent transcription termination prevents misregulation of global gene expression. Nat Microbiol, 1, 15007.

31. Mandell, Z.F., Oshiro, R.T., Yakhnin, A.V., Vishwakarma, R., Kashlev, M., Kearns, D.B. and Babitzke, P. (2021) NusG is an intrinsic transcription termination factor that stimulates motility and coordinates gene expression with NusA. Elife, 10.

32. Dar, D., Shamir, M., Mellin, J.R., Koutero, M., Stern-Ginossar, N., Cossart, P. and Sorek, R. (2016) Term-seq reveals abundant ribo-regulation of antibiotics resistance in bacteria. Science, 352, aad9822.

33. Kosinski, J.G., Ranaweera, S., Chelkowska-Pauszek, A., Kashlev, M., Babitzke, P. and Zywicki, M. (2025) Characterization of bacterial intrinsic transcription terminators identified with TERMITe-a novel method for comprehensive analysis of Term-seq data. Nucleic Acids Res, 53.

34. Brumm, P.J., Monsma, S., Keough, B., Jasinovica, S., Ferguson, E., Schoenfeld, T., Lodes, M. and Mead, D.A. (2015) Complete Genome Sequence of Thermus aquaticus Y51MC23. PLoS One, 10, e0138674.

35. Wang, J., Pan, X. and Liang, X. (2016) Assessment for Melting Temperature Measurement of Nucleic Acid by HRM. J Anal Methods Chem, 2016, 5318935.

36. Kang, J.Y., Mishanina, T.V., Landick, R. and Darst, S.A. (2019) Mechanisms of Transcriptional Pausing in Bacteria. J Mol Biol, 431, 4007–4029.

37. Menendez-Gil, P. and Toledo-Arana, A. (2020) Bacterial 3’UTRs: A Useful Resource in Post-transcriptional Regulation. Front Mol Biosci, 7, 617633.

38. Mitra, A., Angamuthu, K., Jayashree, H.V. and Nagaraja, V. (2009) Occurrence, divergence and evolution of intrinsic terminators across eubacteria. Genomics, 94, 110–116.

39. Cheng, S.W., Lynch, E.C., Leason, K.R., Court, D.L., Shapiro, B.A. and Friedman, D.I. (1991) Functional importance of sequence in the stem-loop of a transcription terminator. Science, 254, 1205–1207.

40. Vargas-Blanco, D.A. and Shell, S.S. (2020) Regulation of mRNA Stability During Bacterial Stress Responses. Front Microbiol, 11, 2111.

41. Blatt, N.B., Osborne, S.E., Cain, R.J. and Glick, G.D. (1993) Conformational studies of hairpin sequences from the ColE1 cruciform. Biochimie, 75, 433–441.

42. Epshtein, V., Cardinale, C.J., Ruckenstein, A.E., Borukhov, S. and Nudler, E. (2007) An allosteric path to transcription termination. Mol Cell, 28, 991–1001.

43. Wilson, K.S. and von Hippel, P.H. (1994) Stability of Escherichia coli transcription complexes near an intrinsic terminator. J Mol Biol, 244, 36–51.

44. Wilson, K.S. and von Hippel, P.H. (1995) Transcription termination at intrinsic terminators: the role of the RNA hairpin. Proc Natl Acad Sci U S A, 92, 8793–8797.

45. Toulokhonov, I. and Landick, R. (2003) The flap domain is required for pause RNA hairpin inhibition of catalysis by RNA polymerase and can modulate intrinsic termination. Mol Cell, 12, 1125–1136.

46. Di Salvo, M., Puccio, S., Peano, C., Lacour, S. and Alifano, P. (2019) RhoTermPredict: an algorithm for predicting Rho-dependent transcription terminators based on Escherichia coli, Bacillus subtilis and Salmonella enterica databases. BMC Bioinformatics, 20, 117.

47. Leinonen, R., Sugawara, H., Shumway, M. and International Nucleotide Sequence Database, C. (2011) The sequence read archive. Nucleic Acids Res, 39, D19–21.

48. Edgar, R.C. (2010) Search and clustering orders of magnitude faster than BLAST. Bioinformatics,26, 2460–2461.

49. Martin, M. (2011) Cutadapt removes adapter sequences from high-throughput sequencing reads. EMBnet.journal, 17.

50. Langmead, B. and Salzberg, S.L. (2012) Fast gapped-read alignment with Bowtie 2. Nat Methods, 9, 357–359.

51. Benson, D.A., Clark, K., Karsch-Mizrachi, I., Lipman, D.J., Ostell, J. and Sayers, E.W. (2015) GenBank. Nucleic Acids Res, 43, D30–35.

52. Li, H., Handsaker, B., Wysoker, A., Fennell, T., Ruan, J., Homer, N., Marth, G., Abecasis, G., Durbin, R. and Genome Project Data Processing, S. (2009) The Sequence Alignment/Map format and SAMtools. Bioinformatics, 25, 2078–2079.

53. Lorenz, R., Bernhart, S.H., Honer Zu Siederdissen, C., Tafer, H., Flamm, C., Stadler, P.F. and Hofacker, I.L. (2011) ViennaRNA Package 2.0. Algorithms Mol Biol, 6, 26.

54. Robinson, M.D., McCarthy, D.J. and Smyth, G.K. (2010) edgeR: a Bioconductor package for differential expression analysis of digital gene expression data. Bioinformatics, 26, 139–140.

55. Yuzenkova, Y., Tadigotla, V.R., Severinov, K. and Zenkin, N. (2011) A new basal promoter element recognized by RNA polymerase core enzyme. Embo J, 30, 3766–3775.

